# A novel role of macrophage PIST protein in regulating *Leishmania major* infection

**DOI:** 10.1101/2023.03.06.531336

**Authors:** Sourav Banerjee, Mandip Pratham Gadpayle, Swagata Das, Suman Samanta, Rupak Datta, Sankar Maiti

## Abstract

PDZ protein interacting specifically with Tc10 or PIST is a mammalian *trans*-Golgi resident protein that regulates subcellular sorting of plasma membrane receptors. PIST has recently been found to play an important role in regulating viral pathogenesis. Alteration in PIST expression is linked to the reprogramming of cell surface receptors which is crucial in determining herpes simplex virus1 infection. In this context, PIST is crucial in triggering autophagy via Beclin 1 -PI3KC3 pathway. However, there is complete lack in our knowledge on the role of this protein in any parasitic infection. *Leishmania* parasites infect their host macrophage cells via phagocytosis and their survival within the parasitophorous compartment has recently been found to be dependent on host autophagy by a yet to be identified mechanism. Using *Leishmania major* (*L. major*)-macrophage infection model system we, for the first-time report here that in infected macrophages Golgi resident PIST protein migrates towards the parasite containing compartment. We have also found that PIST associates with Beclin 1, however, not with LC3 within *L. major* parasite containing compartment of infected macrophages. Further, we performed genetic ablation of PIST by siRNA and observed that knockdown of macrophage PIST in turn helps in parasite replication. Contrary to this, overexpression of PIST in macrophages restricted the multiplication of *L. major*. Collectively, our study for the first time reveals that PIST is essential in regulating intracellular parasite, *L. major* infection within macrophage cells.

**Summary Statement:** Mammalian PIST protein plays a crucial role in regulating cellular trafficking events. Here, we showed that PIST status is altered within *Leishmania major* parasite infected macrophages. Further, we confirmed that PIST is essential in restricting parasite growth. Additionally, a potential interacting axis between PIST and Beclin 1 is identified during infection.

## Introduction

The neglected tropical disease, leishmaniasis is caused by the trypanosomatid group of parasites *Leishmania* that results into ∼1 million new cases every year in about 100 endemic countries worldwide(Burza et al., 2018). *Leishmania* parasites have a digenetic life cycle where they alternate between their sandfly vector and human host (Sunter & Gull, 2017). Once inside its host, the parasite is first phagocytosed by neutrophil cells and further induces apoptosis in these infected cells (Peters et al., 2008). Interestingly, the infected neutrophils are then engulfed by macrophage cells through receptor mediated pathway, without activating macrophage immune responses. This unusual entry process in turn enables the safe delivery of promastigotes into macrophage phagolysosomes (Chang & Dwyer, 1978; Peters et al., 2008). During this process, *Leishmania* loses its flagella and transforms into non-motile amastigotes which continues to replicate within phagolysosomes until the cell bursts (Chang & Dwyer, 1978; Moradin & Descoteaux, 2012). Importantly, this sequential passage from one host cell to another involves dramatic change of environment which possess significant challenges to the parasite. Particularly, while inside the macrophage phagolysosome, *Leishmania* parasites are exposed to oxidative and nitrosative stress, acidic pH. In fact, the nutrient availability within this compartment is also poor (Banerjee & Datta, 2020; Haas, 2007; Thi et al., 2012). However, the parasite has developed several adaptive strategies to overcome these host induced stresses (Burchmore & Barrett, 2001; Chang & Dwyer, 1978; Pal et al., 2017; Späth et al., 2003; van Assche et al., 2011). Although autophagy was originally discovered as a catabolic stress response pathway autonomously regulated by individual cells that degrades or recycles damaged cell contents, increasing evidence already established it to be essential in combating a range of pathogenic infections including *Mycobacterium tuberculosis* (Deretic, 2014). However, activation of autophagic pathway might also be beneficial for the growth of the invading pathogen, e.g., *Staphylococcus aureus* (O’Keeffe et al., 2015; Schnaith et al., 2007). In canonical autophagy process, target molecules or cellular components are separated from the rest of the cell within a double membrane vesicle called autophagosome which is orchestrated by different Atg group of proteins (Mizushima et al., 2011). These proteins further mediate the delivery of cellular cargo containing autophagosomes to lysosome, the centre for degradation of target molecules (Zhao & Zhang, 2019). In this context, it is important to note that differentiation of *Leishmania* parasites within infected macrophages is also dependent on the acidification of parasite containing compartment through fusion with lysosome (ALEXANDER & VICKERMAN, 1975; Zilberstein, 2021). However, if there is any of crosstalk between the phagocytosis process of *Leishmania* parasites and the autophagy machinery in the infected macrophage cells is not clearly known. The earliest report in this field suggested that *L. mexicana* infection induces delivery of nutrients via microautophagy which can then be utilized by the parasite (Schaible et al., 1999). Further studies with different *Leishmania* species confirmed that autophagic induction within macrophage is almost an inevitable outcome of this infection (Cyrino et al., 2012; Dias et al., 2018; Duque et al., 2021; Franco et al., 2017; Pinheiro et al., 2009; Thomas et al., 2018). However, we have a very limited understanding on the underlying mechanism, type of autophagy being induced during infection and its influence on the parasite survival. It has been found that different multiprotein complexes are involved in determining the class of autophagic pathway, including Beclin 1-Vps34 (vacuolar sorting protein-34) complex mediated canonical autophagy or LC3 associated phagocytosis (LAP) (Funderburk et al., 2010; Heckmann & Green, 2019). LAP is a classic example of eliminating invading pathogens which is also known as xenophagy (Bauckman et al., 2015). Recently, Matte *et al* found that *Leishmania* parasites are able to cleave vesicle associated membrane protein 8 (VAMP-8) by the parasite metalloprotease GP63 which in turn inhibits the recruitment of LC3 in parasite containing phagosomes (Matte et al., 2016). Therefore, alteration in LAP during *Leishmania* infection might be beneficial for the parasite to evade the host immune responses. However, what happens to the status of Beclin 1 mediated canonical autophagy during infection remains to be investigated. Interestingly, *Leishmania* infection has been found to promote autophagy in human polymorphonuclear neutrophils (hPMN) that involves phosphorylation of the Beclin 1 protein (Pitale et al., 2019). In addition to this, there was an increased expression of LC3-II protein. Induction of both Beclin-1 and LC3 mediated autophagic pathway were shown to be essential in promoting macrophage engulfment of *Leishmania donovani* infected neutrophils (Pitale et al., 2019). However, this observation contradicts with the *Leishmania* mediated inhibition of LC3 recruitment to parasite containing phagosomes indicating that a different set of events might be happening within the primary host macrophage cells (Matte et al., 2016). Recently, Duque *et al* observed that in *Leishmania braziliensis* infected macrophage cells the transcript level of Beclin1 is upregulated at early time point post infection, however, the status of Beclin 1 protein and its localization in host cells during the parasite infection is yet to be identified (Duque et al., 2021). In this context, it is worth mentioning that based on the environmental que several cofactors or upstream regulators induces autophagy by directing the formation of the Beclin 1-Vps34 core complex which further drives the phagophore construction (Kang et al., 2011). Among these upstream autophagy regulators, PDZ domain protein interacting specifically with Tc-10 (PIST) which is also known as GOPC (Golgi-Associated PDZ and Coiled-Coil Motif-Containing Protein) plays important roles in signal transduction pathway (Ford & Burd, 2022; Koliwer et al., 2015; Neudauer et al., 2001; Wente et al., 2005). Additionally, PIST has been found to be involved in the trafficking of different cellular proteins like Cadherin 23, cystic fibrosis transmembrane conductance regulator (CFTR), and metabotropic glutamate receptor 5 (mGluR5) (Cushing et al., 2008; Klüssendorf et al., 2021; Pelaseyed & Hansson, 2011; Xu et al., 2010). However, a novel role of this protein during pathogenic infection has recently been discovered. It is found that during the Measles virus (MeV) or group A Streptococcus infection (GAS), host epithelial cells activate the complement regulatory protein CD46 and its association with PIST/ GOPC. Further, it was observed that PIST/ GOPC plays a crucial role in inducing autophagy by recruiting the Beclin 1-Vps34 complex (Richetta et al., 2013). Interestingly, silencing of PIST/ GOPC expression failed to induce autophagy during MeV infection (Richetta et al., 2013). However, there is no such report on the status of PIST protein or how it is involved in the Beclin 1-mediated autophagic pathway during any parasitic infection. In this present study, we addressed these unknown issues in *L. major*-macrophage infection model system. We report here that, *L. major* infection induces Beclin 1 protein level at an early time point post infection and localization of this protein in the parasite containing compartment. Importantly, we found that PIST protein is also recruited to *Leishmania* parasite containing phagosome/ phagolysosomal compartment, which is infection specific. We have also observed that during infection, PIST solely colocalizes with Beclin 1 and not with LC3 protein. We have discovered that downregulation/ overexpression of PIST by genetic manipulation leads to significant change in macrophage parasite burden. Our results therefore establishes that PIST is essential in *Leishmania* infection. Collectively, this study for the first time demonstrates the involvement of macrophage PIST protein during *Leishmania* infection by depicting a potential interacting axis with Beclin 1.

## Results

### *L. major* infection induces PIST localization to the parasite containing compartment within the infected macrophages

Previously, it has been shown that PIST protein plays an important role in autophagosome formation during immune signalling (Joubert et al., 2009). However, whether the status/localization of PIST is altered during pathogenic infection is yet to be studied. To address this question, we first sought to investigate the localization pattern of PIST protein in uninfected as well as in *Leishmania* infected macrophage cells systematically. Although it has earlier been reported that in different mammalian cells PIST protein is localized in Golgi compartment we wanted to verify this in macrophages (Lu et al., 2015; Yao et al., 2001). For this, we have first generated a polyclonal antiserum against neuronal form of PIST (Fig. S1A-D) and validated it by western blot in macrophage whole cell lysate (Fig. S1E). As shown in Fig. S2A, in murine J774A.1 cells, our immunofluorescence staining revealed complete colocalization between PIST and Golgin-97 protein confirming its residence solely within the Golgi compartment in resting macrophages which is consistent with the previous reports in the well-known kidney cell lines, COS-7 and MDCK (Lu et al., 2015; Yao et al., 2001). We next, infected J774A.1 macrophages with *L. major* promastigotes for 4- 12hrs and studied if there is any alteration in the status of PIST protein. From our immunofluorescence experiment it was evident that at 4hrs post infection (p.i.), in uninfected macrophages PIST protein is clustered within its native Golgi-compartment (Fig. S2A). Contrary to this, in *L. major* infected macrophages PIST is much more dispersed and present in close proximity of phagocytosed parasites (Fig. 1A). Interestingly, at 6 and 12hrs p.i. with the increase in intracellular parasite burden PIST protein was found to be significantly recruited to the parasite containing compartment (Fig. 1B-C, D). To further validate this observation, we isolated thioglycolate elicited peritoneal macrophages from Balb/c mice and infected them in a similar way. At 6hrs post infection, our immunofluorescence staining recapitulated the same finding where we observed PIST is significantly dispersed from the Golgi compartment in *L. major* infected peritoneal macrophage cells with its simultaneous presence within the parasitophorous vacuole (Fig. S2B). Therefore, our results confirmed *L. major* infection induced alteration of PIST protein in both mouse macrophage cell line as well as in primary macrophages. In fact, our quantitative estimation of PIST positive parasite containing compartment also showed the similar outcome not clear. From Fig. 1E it is evident that at 4hrs p.i. while the percent of PIST positive *Leishmania* containing compartment is ∼60%, it increased upto ∼80% at later time points p.i. However, we did not find any change in PIST protein level throughout the infection time points which is evident from our western blot analysis and its quantification (Fig. 1F, S3). This interesting observation further prompted us to check if this altered localization of macrophage PIST protein is *Leishmania* infection specific.

**Figure 1.**
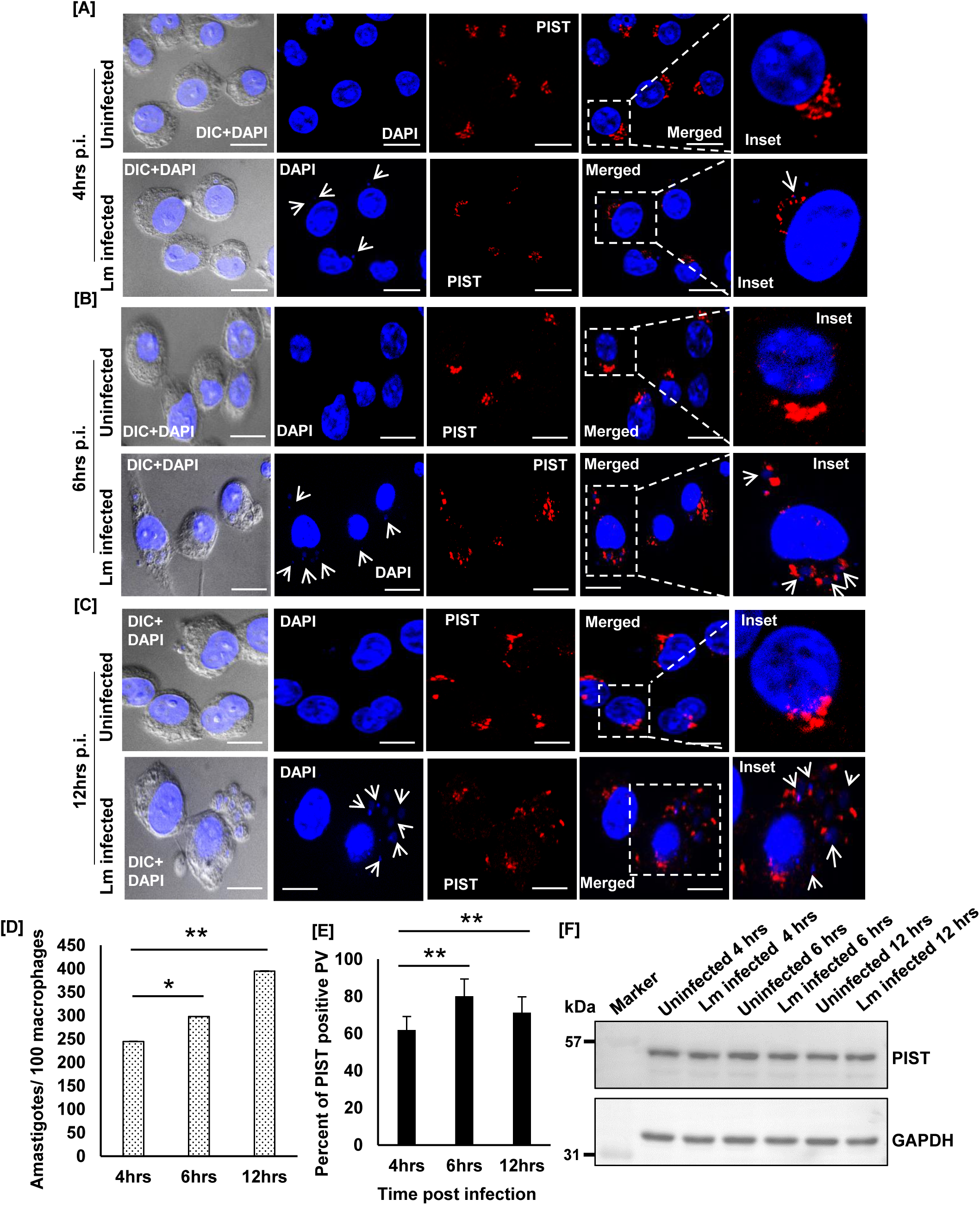
PIST is recruited to the parasite containing compartment of infected macrophages. J774A.1 macrophages were infected with *L. major* (Lm) promastigotes and immunostained with anti-PIST (red) at (A) 4 hrs, (B) 6 hrs and (C) 12 hrs post infection. Nuclei were stained with DAPI (blue). (DIC+DAPI) panel in each time points shows the uninfected and Lm infected macrophage cells where in the DAPI panel the presence of intracellular parasites (smaller nuclei) as indicated by white arrows within infected cells. Insets shows magnified image of a representative cell from both uninfected and Lm infected panels with distribution of PIST protein and arrows indicating parasite nuclei. Cells were visualised with Zeiss Apotome microscope using ×63 oil immersion objective. Scale bar: 20μm. (D) Intracellular parasite burden (amastigotes/100 macrophages) in Lm-infected macrophage cells was measured over 4- 12 hrs p.i. and represented in the bar diagram. (E) Bar diagram represents percent of PIST positive parasitophorous vacuole (PV) at 4- 12 hrs p.i. At least 20 cells from three independent experiments were scored in each of the cases. (F) Cell lysates were prepared from uninfected and Lm infected cells at all the time points post infection and western blot was performed using anti- PIST to verify the protein level of PIST at its predicted molecular weight of ∼60 kDa. GAPDH (∼36 kDa) was used as loading control where the presence of it is shown in the lower panel. Error bars represent mean± standard deviation. **p≤.01; ****p≤.0001

### Altered distribution of PIST protein is *Leishmania* infection specific

To investigate whether disintegration of PIST protein is *L. major* infection specific or not, we had performed latex bead phagocytosis assay. Here, J774A.1 macrophage cells were incubated with polystyrene latex bead (LB) bearing alexa-568 and following its phagocytosis, localization of PIST was observed by immunofluorescence staining. It is interesting to note that in LB treated macrophage cells we could not detect the presence of PIST within LB containing phagosome at 4- 12hrs post incubation (Fig. 2A-C). Also, the distribution pattern of PIST was remained unaltered throughout these time points in both control and LB treated macrophage cells (Fig. 2A-C). This results indeed clearly indicates that recruitment of PIST to the parasite containing compartment was *Leishmania* phagocytosis specific. In this context, it is worth mentioning that we have previously shown that Rab11 is present within parasite containing compartment of *L. major* infected macrophage cells (Banerjee & Datta, 2020). Hence, we further wanted to check whether PIST co-localizes with Rab11 or not while the cells are infected with the parasite. As depicted by our immuno-colocalization experiment in Fig. S4, in *L. major* infected macrophage cells PIST had shown significant colocalization with Rab11 as compared to the uninfected macrophage cells at 12hrs p.i. These together unambiguously demonstrate that PIST was present within parasite containing compartment of infected macrophage cells and the altered distribution of this protein was *L. major* infection specific.

**Figure 2.**
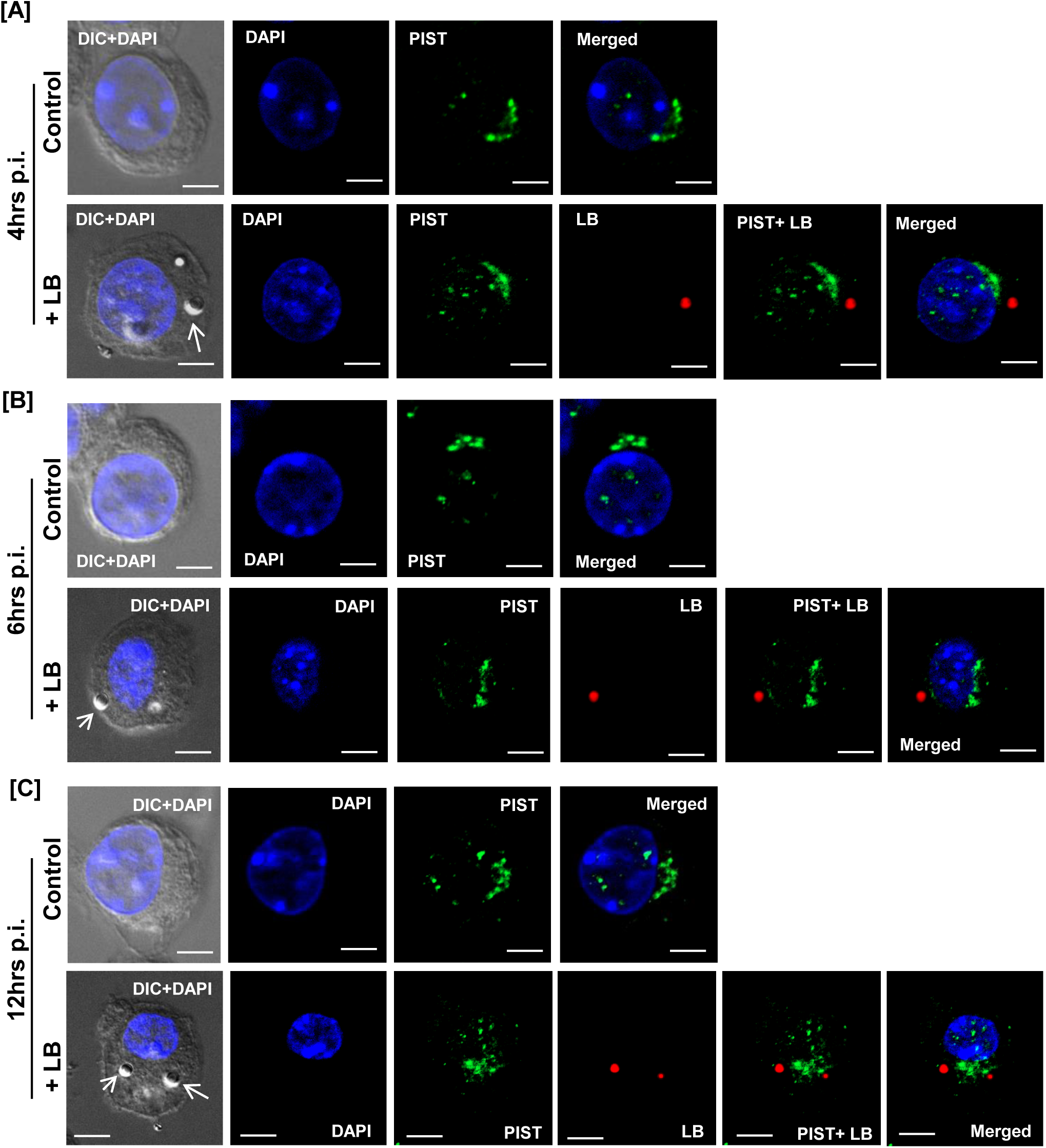
Macrophage PIST localization is unaltered during latex bead phagocytosis. J774A.1 macrophage cells were incubated with FluoSpheres beads (red) at a ratio of 1: 10 (cell: beads) and further immunostained with anti- PIST (green) at (A) 4hrs, (B) 6hrs and (C) 12hrs post incubation. Nuclei were stained with DAPI (blue). (DIC+DAPI) panel in each time points shows the uninfected and latex bead (LB) fed macrophage cells (+LB) where white arrows in (+LB) panel indicative of internalized LB. Cells were visualised with Zeiss Apotome microscope using ×63 oil immersion objective. Scale bar: 20μm.

### *L. major* infection promotes the co-localization of PIST and Beclin 1 within infected macrophages

It has previously been reported that PIST is important in mediating autophagosome formation interacting with VPS34/ Beclin-1 complex (Joubert et al., 2009). Although recent developments demonstrate that different species of *Leishmania* parasites are able to manipulate phagolysosomal biogenesis there is lack in our understanding on the status of Beclin 1 and its role in autophagy during this parasite infection (Cyrino et al., 2012; Dias et al., 2018; Duque et al., 2021; Franco et al., 2017; Pinheiro et al., 2009; Thomas et al., 2018). Therefore, our previous experimental observations prompted us to check if there is any crosstalk between PIST and Beclin 1 during infection. For this, we generated a rabbit polyclonal antiserum against mouse Beclin 1 protein as depicted in Fig. S5. Next, we analysed the status of Beclin 1 in *L. major* infected macrophages for 4- 12hrs p.i. Interestingly, our immunofluorescence microscopic experiment revealed that in infected macrophages Beclin 1 was solely present within *Leishmania* containing compartment (Fig. S6A-C). This further corroborated with our quantitative estimation where percent of Beclin 1 positive parasite containing compartment remained more that 80% throughout the experimental time points (Fig. S6D). Also, our immunofluorescence data showed that the abundance of Beclin 1 within these parasite containing compartments continued to increase at the later time points, i.e., 6 and 12hrs p.i. indicating the formation of the autophagosome (Fig. S6B-C, D). Since, we have already observed the presence of PIST within parasite containing compartment we wanted to check if there is any co-localization between PIST and Beclin 1. Hence, we next analysed colocalization between these two proteins. Although at 4hrs p.i. we did not observe any significant colocalization between PIST and Beclin 1, in *L. major* infected macrophages there was a strong colocalization between these two proteins at 6 and 12hrs p.i. with the MCC of (0.82± 0.1) and (0.81± 0.14) respectively (Fig. 3A-D). However, we were curious to check if this outcome is infection specific or not. Hence, we induced autophagy in macrophages by the well-established method, serum starvation for 6hrs, and further checked for colocalization between PIST and Beclin 1. Interestingly, under such condition, we failed to observe any significant colocalization between these two proteins where the distribution pattern of these two proteins remained similar to the control macrophage cells (Fig. 4). All these together suggests that association of PIST and Beclin 1 is *L. major* infection specific and PIST might play an important role in mediating *Leishmania* containing phagolysosomal maturation by recruiting Beclin 1 in this compartment.

**Figure 3.**
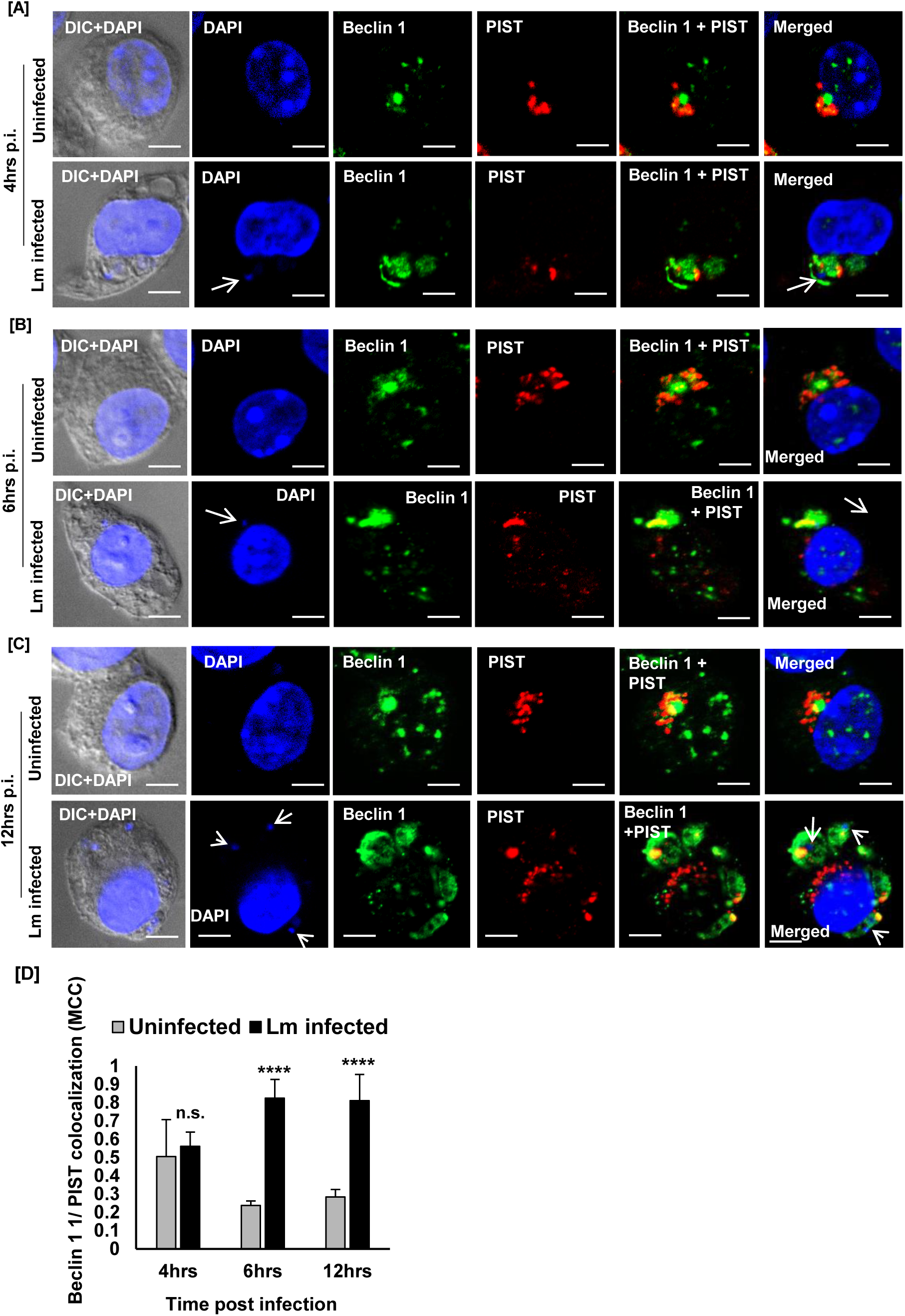
*L. major* infection promotes colocalization of PIST and Beclin 1. J774A.1 macrophages were infected with *L. major* (Lm) promastigotes for (A) 4 hrs, (B) 6 hrs and (C) 12 hrs and further cells were fixed and immunostained with anti-PIST (red) and anti- Beclin-1 (green). Nuclei were stained with DAPI (blue). (DIC+DAPI) panel in each time points shows the uninfected and Lm infected macrophage cells where in the DAPI panel the presence of intracellular parasites (smaller nuclei) is indicated by white arrows. In the merged panel arrows are indicating parasite nuclei. Cells were visualised with Zeiss Apotome microscope using ×63 oil immersion objective. Scale bar: 20μm. (D) Bar diagrams represent Mander’s colocalization coefficient (MCC) of PIST with Beclin 1 measured using ImageJ software. Grey bar represents uninfected macrophages, whereas black bar represents Lm-infected macrophages. At least 20 cells from three independent experiments were scored in each of the cases. Error bars represent mean± standard deviation. ****p≤.0001.

**Figure 4.**
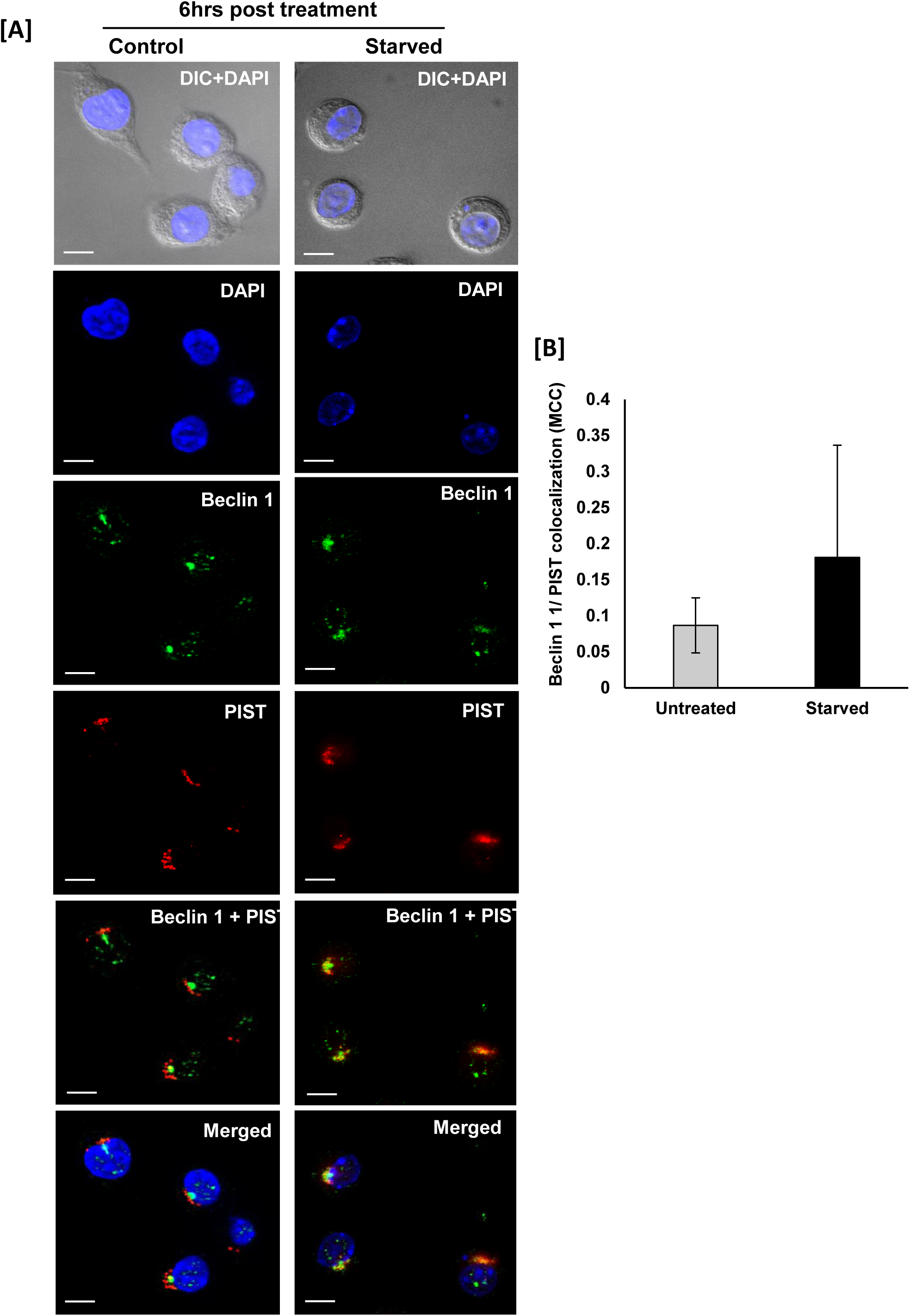
Localization status of PIST and Beclin 1 in serum starved macrophage cells. J774A.1 macrophage cells are initially grown in complete DMEM media and further cells were washed and incubated in serum free medium for 6hrs. (A) Next, cells were fixed and immunostained with anti-PIST (red) and anti- Beclin-1 (green). Nuclei were stained with DAPI (blue). (DIC+DAPI) panel in each time points shows the untreated (control) and starved macrophage cells. Cells were visualised with Zeiss Apotome microscope using ×63 oil immersion objective. Scale bar: 20μm. (B) Bar diagrams represent Mander’s colocalization coefficient (MCC) of PIST with Beclin-1 measured using ImageJ software. Grey bar represents untreated macrophages, whereas black bar represents starved macrophages. At least 20 cells from three independent experiments were scored in each of the cases.

### In *L. major* infected macrophage cells LC3 is not recruited to the PIST containing parasitophorous vacuole

LC3 has recently been found to be involved in phagocytosis via the TLR-signalling pathway that restricts microbial growth (Hayashi et al., 2018). Also, it is important to note that the recruitment of LC3 in Beclin 1 labelled nascent autophagosome is essential for its maturation process (Kang et al., 2011). Hence, taking the clue from our previous observations we next sought to evaluate the involvement of LC3 in the *Leishmania* containing phagosomal compartment. Before proceeding for this, we first checked the status of LC3 protein in *L. major* infected macrophages at 4- 12 hrs p.i. Our immunofluorescence study revealed that in infected macrophage cells LC3 protein is associated in a lesser extent to parasite containing phagosomal compartment where the distribution pattern was remained almost similar with the uninfected macrophages (Fig S7A-C). Previously, Matte *et al*. found that LC3 recruitment to parasite containing compartment in infected macrophage cells is prevented by *L. major* parasites at the early time point (1 hr) post infection (Matte et al., 2016). In this context, we also observed the similar trend. The percent of LC3 positive parasite containing phagosomes remained almost same throughout the experimental time frame of 4- 12hrs (Fig. S7D). This suggests that at late stages post infection LC3 protein recruitment to *Leishmania* containing compartment is largely inhibited as compared to the association with Beclin 1 (Fig. 3 and S6). However, the colocalization status between PIST and LC3 was yet to be determined. Therefore, we further analysed the same systematically at 4- 12hrs p.i. As shown in Fig. 5 our immunofluorescence study failed to find any significant colocalization between PIST and LC3 throughout this time points which might be due to the action of GP63 (Matte et al., 2016). All these together indicates that *L. major* infection inhibits LC3-associated phagocytosis without altering Beclin 1 recruitment. However, it becomes essential to identify the role of PIST during *Leishmania* infection.

**Figure 5.**
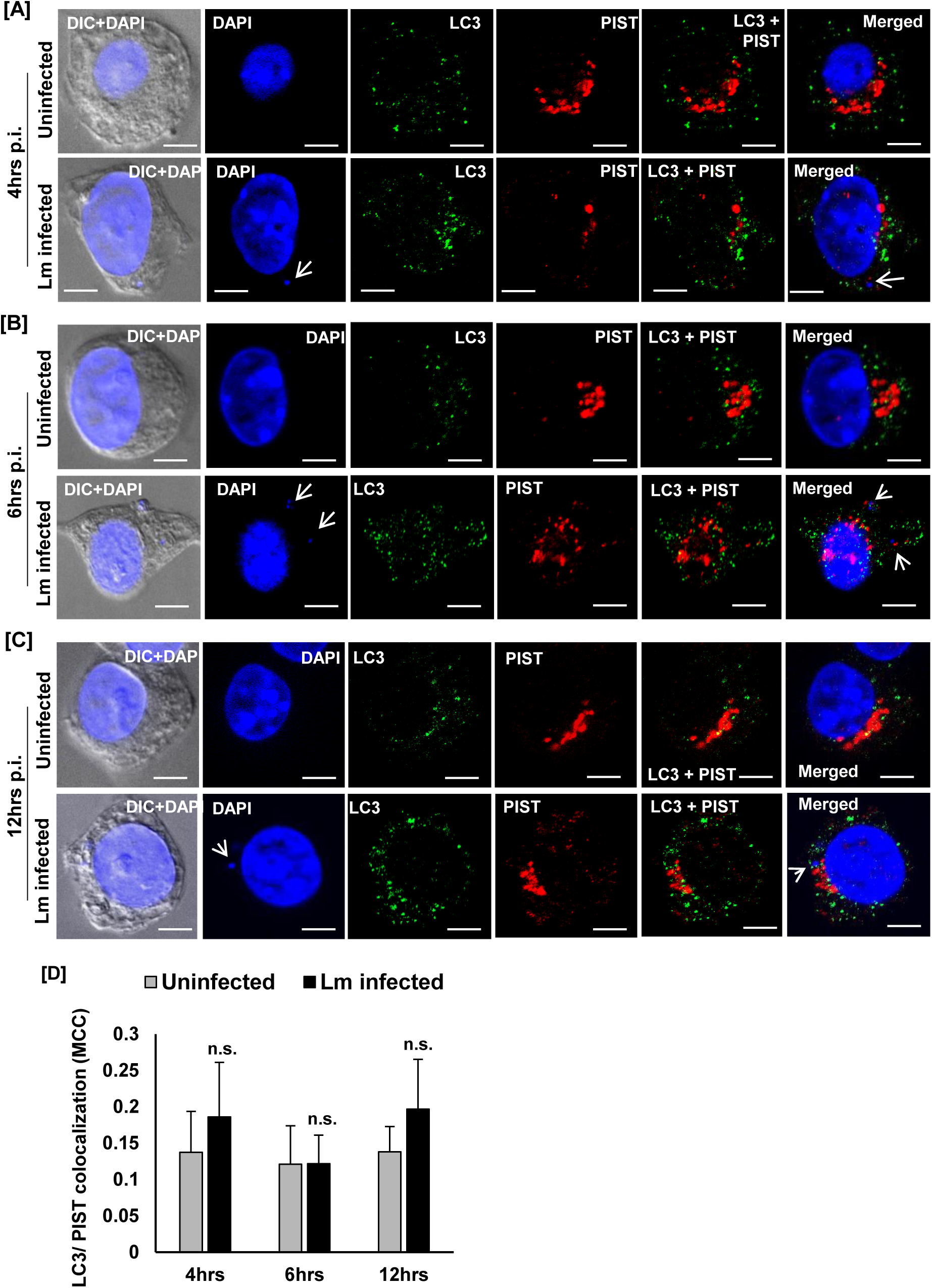
LC3 is not recruited to the PIST positive parasitophorous compartment in *L. major* infected macrophages. J774A.1 macrophages were either uninfected or infected with *L. major* (Lm) promastigotes for (A) 4 hrs, (B) 6 hrs and (C) 12 hrs and further cells were fixed and immunostained with anti-PIST (red) and anti- LC3 (green). Macrophage and parasite nuclei were stained with DAPI (blue). (DIC+DAPI) panel in 4- 12 hrs time points shows the uninfected and Lm infected macrophage cells. Arrows in DAPI and ‘Merged’ panel are indicative of intracellular parasites (smaller nuclei). Cells were visualised with Zeiss Apotome microscope using ×63 oil immersion objective. Scale bar: 20μm. (D) Bar diagram represents Mander’s colocalization coefficient (MCC) of PIST with LC3 protein in uninfected (grey bar) and Lm infected (black bar) measured using ImageJ software. At least 20 cells from three independent experiments were scored in each of the cases. Error bars represent mean± standard deviation. n.s. non-significant.

### Macrophage PIST is essential in controlling *L. major* infection

Our experimental results suggests that PIST might be playing a crucial role in the maturation process of *Leishmania* containing phagosomes by recruiting Beclin 1 in infected macrophage cells. However, the unavailability of information on the function of PIST during pathogenic infection possesses a major limitation in our understanding. Therefore, to evaluate the function of PIST during *L. major* infection we next attempted to overexpress this protein and check its effect on infectivity. For this, we have transiently overexpressed nPIST (neuronal form) in J774A.1 macrophage cells. These cells were further infected with *L. major* promastigotes and the parasite burden was measured at 6hrs p.i. In the overexpressing cells heightened level of PIST protein was found to be clustered within amastigote containing phagosomal compartment (Fig. 6A). Interestingly, in these cells the intracellular parasite burden was significantly reduced as compared to the untreated infected control macrophages (Fig. 6C). This result strongly suggests that macrophage PIST is crucial in augmenting *Leishmania* infection. However, we further wanted to validate what happens to the parasite infectivity if PIST is downregulated. For this, we generated macrophage cells where PIST expression is suppressed by siRNA. These cells were further infected with *L. major* promastigotes similarly and the amastigote/ macrophage count was measured at 6hrs p.i. Importantly, as compared to the control, in these cells expressing reduced PIST protein, we found significantly higher parasite burden compared to the untreated infected macrophages (Fig. 6C). All these together unambiguously confirmed for the first time that PIST protein is essential in controlling *L. major* infection in macrophage cells.

**Figure 6.**
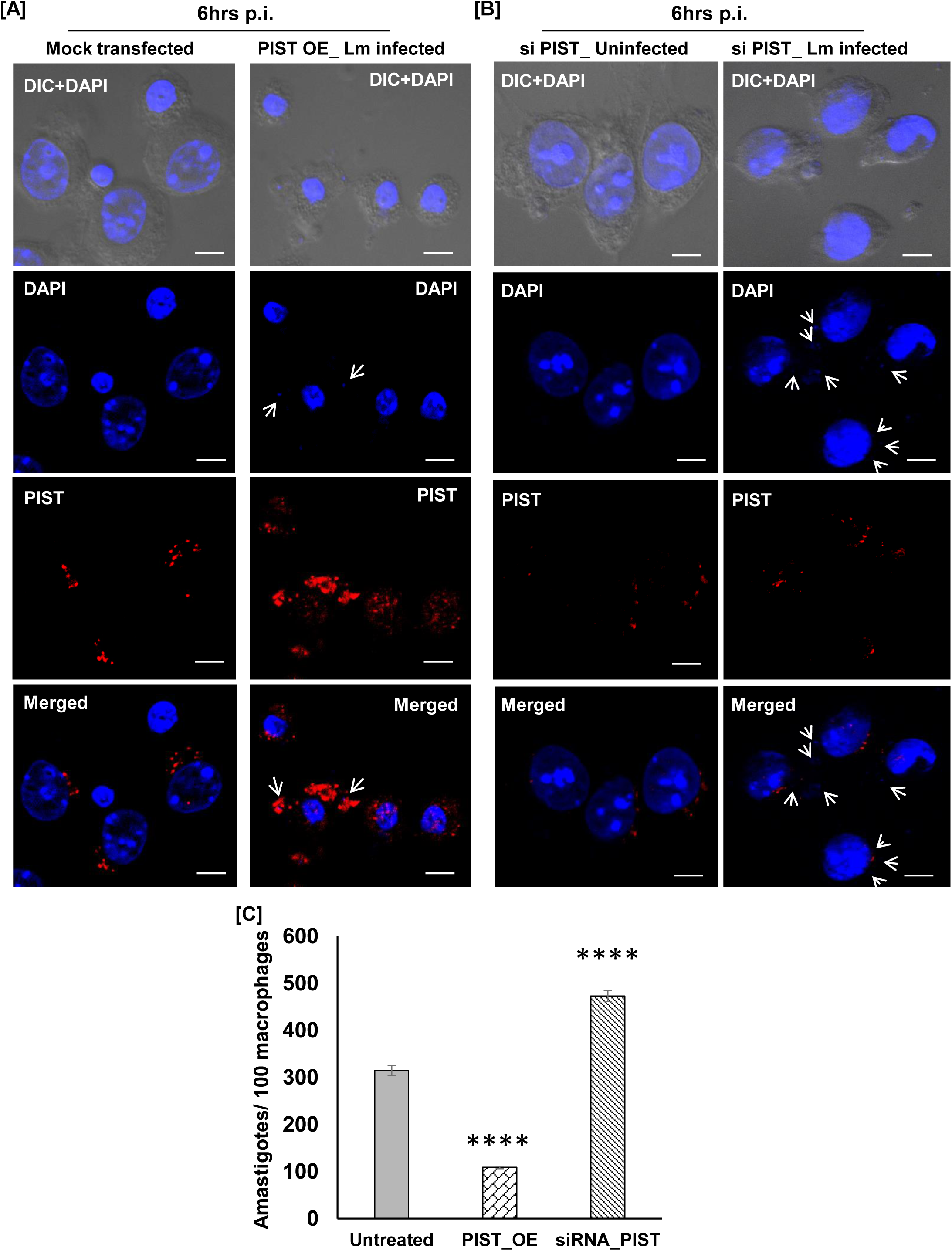
PIST is essential in controlling *L. major* infection in infected macrophage cells. (A) J774A.1 macrophage cells were either mock transfected (vector control) or transfected with pEGFPC1_PIST to overexpress PIST protein as described in the materials methods. (B) Contrary to this, to knockdown the expression of PIST, macrophage cells were transfected with siRNA against human GOPC. Now (A) the overexpressing cells or (B) siRNA knockdown cells were infected with *L. major* (Lm) promastigotes for 6 hrs following which cells were fixed and immunostained with anti-PIST (red) to check the status of this protein. In both the cases either (A) mock transfected (vector control) or the (B) siRNA knockdown cells were kept uninfected and used as control. nPIST_OE Lm infected panel is representative of overall PIST protein level including endogenous expression. Macrophage and parasite nuclei were stained with DAPI (blue). (DIC+DAPI) panel shows either control or Lm infected cells. Arrows in DAPI and ‘Merged’ panel are indicative of intracellular parasites (smaller nuclei) within Lm infected macrophage cells. Cells were visualised with Zeiss Apotome microscope using ×63 oil immersion objective. Scale bar: 20μm. (C) Intracellular parasite burden in either PIST overexpressed or in PIST siRNA knockdown Lm infected macrophage cells were measured 6 hrs p.i. Intracellular parasite burden (amastigotes/100macrophages) was quantified from at least 100 macrophage cells. Error bars represent mean± standard deviation values calculated from at least three independent experiments. **p≤.01; ****p≤.0001.

## Discussion

Autophagy is an essential metabolic process that enables cells to autonomously digest and degrade damaged or unnecessary contents (Mizushima et al., 2008). There are a diverse range of autophagy processes including macroautophagy, microautophagy, chaperon mediated autophagy and non-canonical autophagy that enables the digestion of target molecule within lysosomes (Codogno et al., 2012; Parzych & Klionsky, 2014). Although its role in recycling of cellular metabolites has long been established recent investigations has revealed that autophagy is potentially a primordial form of eukaryotic immunity against invading pathogens including *Mycobacterium tuberculosis*, *Salmonella typhi* (Deretic, 2014; O’Keeffe et al., 2015; Schnaith et al., 2007). In case of the protozoan parasite *Leishmania* infection, induction of host autophagy has been found to be an essential outcome (Cyrino et al., 2012; Dias et al., 2018; Duque et al., 2021; Franco et al., 2017; Pinheiro et al., 2009; Thomas et al., 2018). However, the underlying mechanism behind the autophagy induction during *Leishmania* infection is not clear. The non-canonical LC3-associated autophagy plays a crucial role in the clearance of different pathogenic infections like *L. monocytogenes*, *Aspergillus fumigatus,* and *Saccharomyces cerevisiae* by macrophage cells (Akoumianaki et al., 2016; Lamprinaki et al., 2017; Li et al., 2021). However, this process has recently been found to be inhibited by *Leishmania* parasite (Matte et al., 2016). Therefore, it becomes essential to study whether the canonical autophagy process is also disrupted during *Leishmania* infection. Beclin 1 is a Bcl-2 homology (BH)-3 domain only protein that determines the progression of autophagy by regulating autophagosome formation and its maturation (Funderburk et al., 2010). This autophagosome biogenesis process is dependent on the functional Beclin 1-Vps34-Vps15 core complex which is mediated by multiple interacting partners of Beclin 1 (Funderburk et al., 2010). Among these proteins, different pathogens ranging from virus to bacteria, have found to alter the functioning of GOPC/PIST protein in order to disrupt the canonical autophagy induction which could be potentially through Beclin 1 as PIST has been established to be an interacting partner Beclin 1 (Joubert et al., 2009). In this background, we have attempted for the first time to identify the status of macrophage Beclin 1-PIST axis during *Leishmania* infection.

Previously, it was reported that *L. braziliensis* infection upregulates Beclin 1 transcript level at early time point post infection, however, as of now there is no information available on what happens to this protein during the course of *Leishmania* infection (Duque et al., 2021). In this context, our immunofluorescence experiment revealed that Beclin 1 protein level remains significantly high in *L. major* infected J774A.1 macrophages throughout the experimental time points, 4- 12 hrs p.i. Importantly, compared to uninfected control cells, in infected macrophages, Beclin 1 was found to be present mostly within the parasite containing compartment. Although Pitale *et al*. has recently shown that *L. donovani* infection promotes interaction between Beclin 1 and Rubicon protein, it seems that there was not any significant alteration in the distribution of Beclin 1 (Pitale et al., 2019). Also, the information available on this aspect is mostly at early time points post infection. However, our temporal studies had shown that localization of Beclin 1 changes upon infection as early as 4hrs p.i. and it remains associated with *Leishmania* containing compartment even at late time point, e.g., 12hrs p.i. It may appear to be inconsistent with the previous report, however, previous studies were performed on hPMNs, whereas we conducted the experiments on macrophage cells, the primary host of *Leishmania* parasites. Therefore, this observation clearly indicates that in macrophage cells *Leishmania* infection might involve different set of signalling cascade. In this context, it is worth mentioning that in *Toxoplasma gondii* infection, Beclin 1 is found to play an important role against the parasite infection (Portillo et al., 2010; Wu et al., 2020). However, the alteration of Beclin 1 protein status along with its localization in parasite containing compartment throughout *Leishmania* infection in infected macrophages is certainly a new finding. Previously, respiratory syncytial virus (RSV) was found to induce Beclin 1 protein level by inhibiting its degradation to drive enhanced autophagy (Chiok et al., 2022). In fact, increased phosphorylation of Beclin 1 at early time point has been reported during *L. donovani* infection (Pitale et al., 2019). Therefore, we speculate it is possible that *Leishmania* infection promotes increased availability of active Beclin 1 protein in macrophage cells. Interestingly, we did not observe any change in LC3 protein abundance in *Leishmania* infected macrophage cells. Also, we did not observe any significant recruitment of LC3 protein in the parasite containing compartment even at late stages post infection as the previous report (Matte et al., 2016). Collectively, these results support the notion that *L. major* infection does not trigger LAP, however, we could not rule out the possibility of autophagy induction via Beclin 1 dependent pathway. Importantly, it would be highly rewarding to check if any interacting partner protein of Beclin 1 is involved in this process. Other than its essential role in protein targeting, PIST has been found to be a potential binding partner of Beclin 1 (Joubert et al., 2009). Importantly, the interaction between Beclin 1-PIST is established to be essential in inducing autophagy. Although there are few other binding partners of Beclin 1, the involvement of PIST specifically during pathogenic infections is certainly a novel aspect (Joubert et al., 2009). During MeV or Group A *Streptococcus* infection the complement activation regulatory protein CD-46 upon recognizing the pathogen signals GOPC/PIST which then mediates the recruitment of Beclin 1-Vps34 complex (Joubert et al., 2009; Richetta et al., 2013). This cross-linking of the CD46-Beclin 1-Vps34 complex via GOPC/PIST is essential in formation of autophagosome and further autophagy process during MeV infection (Richetta et al., 2013). In this background, since we have also noticed a significant trafficking of Beclin 1 to the *Leishmania* containing compartment, it also indicated that macrophage PIST protein might have a crucial role in this process. In this context, our immunofluorescence results demonstrated that upon *L. major* infection, macrophage PIST localization is altered significantly. While in control macrophage cells PIST was concentrated in trans-Golgi region /or TGN, with increasing time points post infection, localization of PIST is completely distorted. Most interestingly, PIST was found to be present in the parasite containing compartments. So far, there is no such attempt to check modulation of PIST localization during infection, hence, we decided to further validate this result. Interestingly, in our latex bead phagocytosis assay under the same experimental set up we failed to observe any discrepancy in localization of PIST at 4- 12hrs post incubation which unequivocally proved the process to be infection specific. Therefore, our data for the first time showed movement of PIST to the parasitophorous vacuole within infected macrophage cells. However, if PIST physically interacts with Beclin 1 or not is yet to be checked. In this context, our immuno-localization study revealed that in uninfected macrophage cells PIST is mostly present surrounding the Beclin 1 protein, which is likely because both these proteins are primarily found in the Golgi network (Kihara et al., 2001; Lu et al., 2015; Yao et al., 2001). Contrast to this, in infected macrophages PIST was found to colocalize completely with Beclin 1 where both of these two proteins are present in parasite containing compartment. Importantly, in starved macrophage cells we did not observe similar fate where the colocalization pattern were comparable with the control cells. In fact, we also failed to detect any colocalization between LC3 and PIST during *L. major* infection. Therefore, based on the previous reports as well as our current observations, it seems that PIST might play a crucial role in the recruitment process of Beclin 1 to the *Leishmania* parasitophorous vacuole similar to its role during MeV infection (Richetta et al., 2013). However, if there is any upstream protein factor which may induce this trafficking process during *Leishmania* infection remains to be an open area of research.

It is interesting to note that herpes simplex virus-1 (HSV-1) has recently been found to promote degradation of GOPC/PIST in order to escape cellular recognition pathway (Soh et al., 2020). It was also shown that toll like-receptor 2 (TLR-2) functioning is dependent on the GOPC/ PIST protein suggesting that in addition to regulating autophagy PIST plays an important role in activating cellular defense against pathogens (Soh et al., 2020). Since, our data shows that upon *L. major* infection, macrophage PIST protein is significantly modulated we further wanted to check if PIST is important in controlling the parasite infection. Therefore, to acquire a complete knowledge on this aspect, we attempted to perform both overexpression as well as knockdown of PIST in macrophage cells and analysed its impact over *Leishmania* infection. We could successfully overexpress this protein in J774A.1 macrophage cell line as evident from our immunostaining experiment. While the PIST overexpressing cells were infected with *L. major* promastigotes we observed a marked increase in the level of PIST protein surrounding the parasite containing compartment. Interestingly, it had a drastic effect over *Leishmania* infection as evidenced by the significant decrease of intracellular parasite burden at 6hrs p.i. This result was further corroborated with our PIST knockdown experiment. In siRNA mediated PIST knockdown macrophage cells, the parasite burden was strikingly high compared to the control infected macrophage cells. Since, *Leishmania* parasites are able to bypass the LC3 associated phagocytosis, based on our observations, it seems that to counteract this event host is trying to induce Beclin 1 mediated autophagy via PIST protein (Matte et al., 2016). Taken together, these results confirmed that macrophage PIST is crucial in controlling *Leishmania* infection. However, the underlying mechanism behind this would certainly be a hot topic of research in future. Also, it would be interesting to see how the early association of PIST with the *Leishmania* parasite containing compartment effect its maturation process. Nevertheless, our study definitely provided a significant knowledge for the first time on the role of PIST in regulating an intracellular parasite infection using macrophage- *L. major* infection model system.

## Materials & Methods

All reagents were purchased from Sigma-Aldrich unless mentioned specifically. Primers used for PCR were obtained from Integrated DNA Technologies.

### *Leishmania* and mammalian Cell culture

The *L. major* strain 5ASKH was a kind gift of Dr. Subrata Adak (IICB, Kolkata). Briefly, *L. major* promastigotes were cultured in M199 medium (Gibco) pH 7.2, supplemented with 15% heat-inactivated fetal bovine serum (FBS, Gibco), 23.5 mM HEPES, 0.2mM adenine, 150 µg/ml folic acid, 10 µg/ml hemin, 120 U/ml penicillin, 120 µg/ml streptomycin, and 60 µg/ml gentamicin at 26°C. J774A.1 murine macrophage cells (obtained from National Center for Cell Sciences, Pune) were grown in Dulbecco’s modified Eagle’s medium (DMEM, Gibco) pH 7.4 supplemented with 2 mM L- glutamine, 100 U/ml penicillin, 100 μg/ml streptomycin, and 10% heat-inactivated FBS at 37°C in a humidified atmosphere containing 5% CO2. Cell number was quantified using a hemocytometer as reported by us previously (Banerjee & Datta, 2020).

### Isolation of peritoneal macrophages from BALB/c mice

BALB/c mice were obtained from the National Institute of Nutrition (NIN), Hyderabad, and they were housed in the institutional animal facility. The CPCSEA guidelines and the Institutional Animal Ethics Committee-approved protocol were followed to conduct experiments with these mice. From 6–8 weeks old mice thioglycolate-elicited peritoneal macrophages were isolated as described earlier (Banerjee & Datta, 2020; Zhang et al., 2008). Briefly, 4 days after intraperitoneal injection of 3% Brewer’s thioglycolate medium (Himedia), peritoneal macrophages were collected using 20G needle from euthanized mice. The isolated macrophages were cultured in complete DMEM (pH 7.4) with 10% heat inactivated FBS at 37°C in a humidified atmosphere containing 5% CO2. Non-adherent cells were discarded after 18hrs. Cellular viability was determined using trypan blue dye exclusion test.

### Infection of macrophage cells, latex bead phagocytosis and starvation assay

Either primary peritoneal macrophage or J774A.1 macrophage cells were infected following the same protocol as described by us previously (Banerjee & Datta, 2020). Briefly, J774A.1 murine macrophages were infected with stationary phase *L. major* promastigotes or for phagocytosis assay, incubated with FluoSpheres Fluorescent Microspheres (580/ 605) (Molecular Probes, Life Technologies) (Kind gift of Dr. Bidisha Sinha, IISER Kolkata) at a ratio of 1: 10 macrophage: parasite/ bead for either 4hrs, 6hrs or 12hrs depending on the experiments. Following infection, cells were washed, fixed with acetone-methanol (1:1) and mounted with anti-fade mounting medium containing DAPI (VectaShield from Vector Laboratories) to visualize the macrophage and parasite nuclei. The phagocytosis of latex beads was confirmed by observing fluorescence emission at red channel. Quantification of intracellular parasite burden (number of amastigotes/100 macrophages) was performed by counting the total number of DAPI-stained nuclei of macrophages and *L. major* amastigotes in a field (at least 100 macrophages were counted from triplicate experiments). For the starvation assay, macrophage cells grown on glass coverslips in normal DMEM media were washed with sterile PBS. Next, the cells were incubated with serum free DMEM media for 6hrs. Following this, cells were processed for immunofluorescence experiment.

### Immunofluorescence

J774A.1 macrophage cells grown on glass coverslips were fixed with acetone: methanol (1:1) at room temperature for 10 min. Further, cells were permeabilised using 0.1% triton-X 100 and blocked with 0.2% gelatin for 5 min at room temperature. Cells were then incubated with desired primary antibodies anti-PIST 1:100; anti-Beclin 1 1:200 and anti-LC3 (CST- rabbit monoclonal against LC3B) 1:50; anti- Rab11 antibody (Santa-Cruz Biotechnology) 1:100 for 1.5 hrs at room temperature and thereafter washed with PBS. Cells were then incubated with secondary antibodies (molecular probes), either goat anti-rabbit Alexa fluor 488 (1:800), or goat anti-mouse Alexa fluor 568 (1:600) and washed with PBS. Finally, the cells were mounted on antifade mounting medium containing DAPI and imaged in Carl Zeiss Apotome.2 microscope using ×63 and ×100 oil immersion objectives or in Olympus IX-81 epifluorescence microscope to monitor parasite burden using ×60 objectives. In colocalization experiments, Pearson’s correlation coefficient (PCC) was calculated as described previously (McDonald & Dunn, 2013). Relative fluorescence intensity was measured using microscope’s own software ZEN Blue as described previously.

### Overexpression of nPIST in macrophage cells

J774A. 1 Macrophage cells were grown till their confluency reached 80-90%. The growth media in these cells was replaced by 1 ml serum deficient DMEM. Meanwhile, the transfection mixture for each plate was prepared. Both the DNA (pEGFPC1-nPIST) (8μg) and lipofectamine-2000 at a ratio of 1:2.5 (w/v) were diluted in serum deficient DMEM separately and incubated at RT for 5min. Thereafter both these suspensions were combined, mixed gently, and incubated again at RT for 20 min. This DNA-lipofectamine mix was then added into the confluent macrophage cell plates containing 1 ml serum reduced DMEM and kept in incubation for 4hrs. After that transfection mix was replaced by selection media containing antibiotic G418 (50ug/ml) in normal cell growth media. Transfected cells were selected in this selection media for the next two days following which the selection media was removed. To infect the cells with *L. major*, same process was followed as described earlier.

### Knockdown of PIST with siRNA

To perform the siRNA mediated knockdown of macrophage PIST, the DharmaFECT Transfection Reagents were used following the manufacturers (horizon, UK) siRNA transfection protocol using the siRNA against siGENOME Human GOPC (57120) siRNA-SMARTpool (#M-020464-01-0005, Dharmacon) and siGENOME non-Targetting siRNA Pool, #1 (#D-001206-13-05, Dharmacon) (Reynolds et al., 2004). Briefly, 5 μM siRNA solution in 1X siRNA buffer was prepared. Now the siRNA is diluted with serum free medium and the Lipofectamine 2000 reagent as well separately according to the manufacturer’s instructions. Now these two solutions are mixed properly and incubated at room temperature for 20 min. The culture medium from J774A.1 macrophage cells growing in 6 well plate was removed and the siRNA containing transfection medium added to each well and kept in incubation at 37°C in 5% CO2 for 6 hrs. Following this, the cells were washed twice with serum free DMEM medium and further complete DMEM added (with 10% FBS) and kept in incubation for 12 hrs. For the infection studies, *L. major* promastigotes at a ratio of 1: 30 (macrophage: parasite) now added to the cells and performed the experiments for desired time point.

### Plasmid Construction

Mouse nPIST was cloned into bacterial expression vector pET28a had been used for N-terminal histidine-tag protein purification. nPIST gene subcloned into pEGFP-C1 mammalian expression vector used for the transient expression. All the mentioned nPIST clones were prepared in our previous preprint (Das et al., 2018 *Preprint*). Total RNA from the C57BL/6 mouse brain was extracted using TRIzol (Invitrogen) from the adult mouse brain tissue with the manufacturer’s protocol. From extracted total RNA, cDNA was prepared by RT PCR kit (Superscript III, Invitrogen), and BECN 1 (1^st^-445^th^ amino acid) gene was amplified using forward primer (5’- GCGGGATCCATGGAGGGGTCTAAGGCGTC-3’) and reverse primers (5’- CGCAAGCTTTCAGAACTGTGAGGACACCCAGGC-3’). Mouse BECN 1/Beclin 1 gene cloned into a pEGFP-C1 mammalian expression vector using BglII and HindIII and confirmed by sequencing. The Beclin 1, 1-268 fragment was subcloned into a modified pET29b (Novagen) bacterial expression vector using forward primer (5’-GCGGGATCCATGGAGGGGTCTAAGGCGTC-3’) and reverse primer (5’-CGCGAAGCTTGAAGACATTGGTTTTCTTGAGC-3’) for C-terminal histidine-tag protein purification.

### Protein Purification and Immunization

nPIST full length and Beclin-1 fragment 1-268 construct transformed into *E. coli* BL-21 DE3 strain (Stratagene) for protein expression and purification purposes. Transformed *E. coli* cells with both constructs were grown in the presence of kanamycin (50µg/ml, Himedia) for 8-10 hrs. 1% inoculum was taken from 8-10 hrs grown primary culture, and the given secondary culture was further grown up to 0.6 Optical Density at 37°C temperature with 150 rpm rotation. After reaching 0.6 O. D culture was induced by 0.5 mM IPTG (Himedia) and further grown for 4hrs at 37°C (nPIST) and for 8hrs at 25°C (Beclin-1 fragment 1-268), harvested bacterial pellets were stored in -80°C (Moseley et al., 2006). nPIST-containing bacterial cells were further processed and protein purified by following the protocols used in the preprint (Das et al., 2018 Preprint). The bacterial cell containing the construct of Beclin-1 fragment 1-268 was re-suspended in ice-cold lysis buffer (50 mM Tris-Cl pH-8.0, 100 mM NaCl, 30 mM Imidazole pH 8.0, 0.5% IGEPAL (Sigma-Aldrich), 0.5 mM DTT, 0.2% sarcosine, 0.2% Thesit (Sigma-Aldrich), 2.5 M guanidine hydrochloride (Sigma-Aldrich) and 1X protease inhibitor cocktail) and subjected to lysis by sonication (Vibra Sonics) for 6 minutes with 35% amplitude, 30 seconds pulse and 30 seconds intervals. The semi-transparent lysed cell solution was centrifuged at 12500 rpm (4°C) for 15 minutes. The supernatant was collected in a fresh tube and further incubated with Ni- NTA resin beads (Qiagen) for 2 hr at 4°C with 5 rpm rotation. After incubation, Ni-NTA beads were washed with wash buffer (50 mM Tris-Cl pH 8.0, 100 mM NaCl, and 30mM Imidazole pH 8.0), and elution was done with elution buffer (50 mM Tris-Cl pH 8.0, 150 mM NaCl, 350 mM Imidazole pH 8.0 and 5% glycerol). The eluted proteins sample were further dialyzed in HENG_5_ buffer (20 mM HEPES, 1 mM EDTA, 150 mM NaCl, and 5% glycerol) for 8 hrs at 4°C temperature. After dialysis, the purified protein samples of nPIST and Beclin 1 fragment 1-268 were stored at 4°C.

Purified recombinant protein nPIST and Beclin 1 were used for immunization and antisera generation in mice and rabbits respectively. All the antisera were raised by following the Institutional Animal Ethics Committee-approved protocols, and CPCSEA, Govt. of India guidelines were followed strictly during conducting the animal experiments.

### Western blot analysis

To check the protein level in uninfected and *L. major* infected macrophage cells J774A.1, cells were scrapped, washed, and lysed in a lysis buffer containing 50mM Tris-Cl (pH 7.5), 150mM NaCl, 0.5% SDS, 10mM NaF, 2mM EDTA, 1mM PMSF, and 1X protease inhibitor cocktail (PIC). Lysis was done using a small probe sonicator at 100 amplitude, 5 seconds on and 30 seconds off. All the process of lysis was strictly performed on ice. After lysis, total protein estimation was done using the filter paper dye-binding method (Minamide & Bamburg, 1990), 10 µg of total protein was loaded in each well, and the proteins were resolved by 10% SDS-PAGE. The resolved gel was subjected to transfer on a PVDF membrane (0.45µM) using semi-dry transfer apparatus (BIO-RAD) after the transfer blot was further blocked in a blocking buffer containing 5% skim milk for 4 hours. The blot was incubated with primary Anti-PIST polyclonal antibody-containing serum (1:1000) and Anti-GAPDH antibody (1:4000, Bio Bharati) in 1X TBS overnight at 4°C, washed with 1X TBST (0.2% Tween-20) and further incubated with goat anti-mouse IgG (Thermo Scientific) and goat anti-rabbit (Thermo Scientific) secondary antibodies conjugated with horse radish peroxidase (HRP) for 2 hours at room temperature. The blot was finally washed with 1X TBST and developed using SuperSignal West Pico Chemiluminescence substrate (Thermo Scientific), and the blot images were captured using the SYNGENE G: BOX, Chemi XRQ.

## Acknowledgments

The authors would like to thank Dr. Priyanka Datta (IISER Pune, India) for providing the mouse Beclin 1 and nPIST construct. S. B. was supported by DST-INSPIRE PhD fellowship and M.P.G. is supported by the CSIR-PhD fellowship. S.S. and S.D. are supported by UGC and IISER Kolkata PhD fellowships respectively. The authors sincerely thank Mr Ritabrata Ghosh for his expert technical assistance.

## Author Contributions

S.B., M.P.G., SS, S.D., and S.M. designed experiments. S.B. and M.P.G. performed the experiments and data analysis. S.B., M.P.G., S.M., and R.D. wrote the manuscript.

---

**Figure S1:**
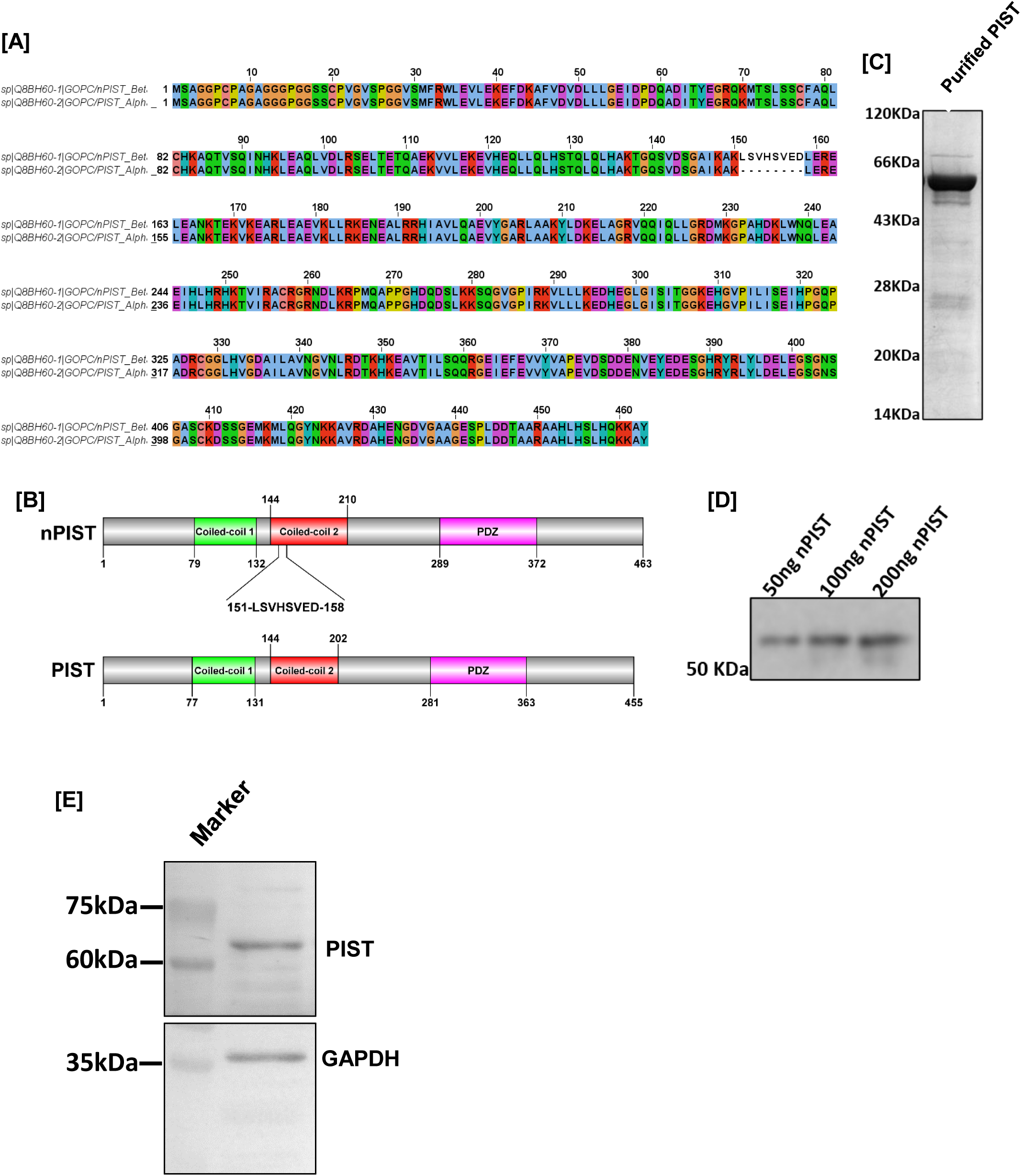
Generation of polyclonal antiserum against nPIST. A. Sequence alignment of neuronal PIST with the non-neuronal form of PIST/GOPC (UniProt accession no. Q8BH60) was performed using Clustal Omega and the visualization file was generated using Jalview. B. Schematic diagram of full-length nPIST (neuronal form) and full-length PIST, both contain two coiled-coil domains (CC1 and CC2) and a PDZ domain, the neuronal form of PIST contains extra 8 amino acids in the Coiled-coil 2 domain. The structure of nPIST/PIST and domains were created using software DOG2.0 (Ren et al., 2009). C. Purified N-terminal Histidine tagged nPIST expressed into *E. coli* BL-21 DE3 strain used to raise mouse anti-nPIST polyclonal antibody; D. Western blot analysis was performed with 1:1000 dilution of immunized mouse serum against the recombinant nPIST (50ng, 100ng, and 200ng). (E) 10 µg of J774A.1 macrophage whole cell lysate was subjected to western blot by anti PIST (1:500) and using anti-GAPDH (1:3000) as loading control. The appearance of PIST protein is indicated at its predicated molecular weight ∼60 kDa.

**Figure S2.**
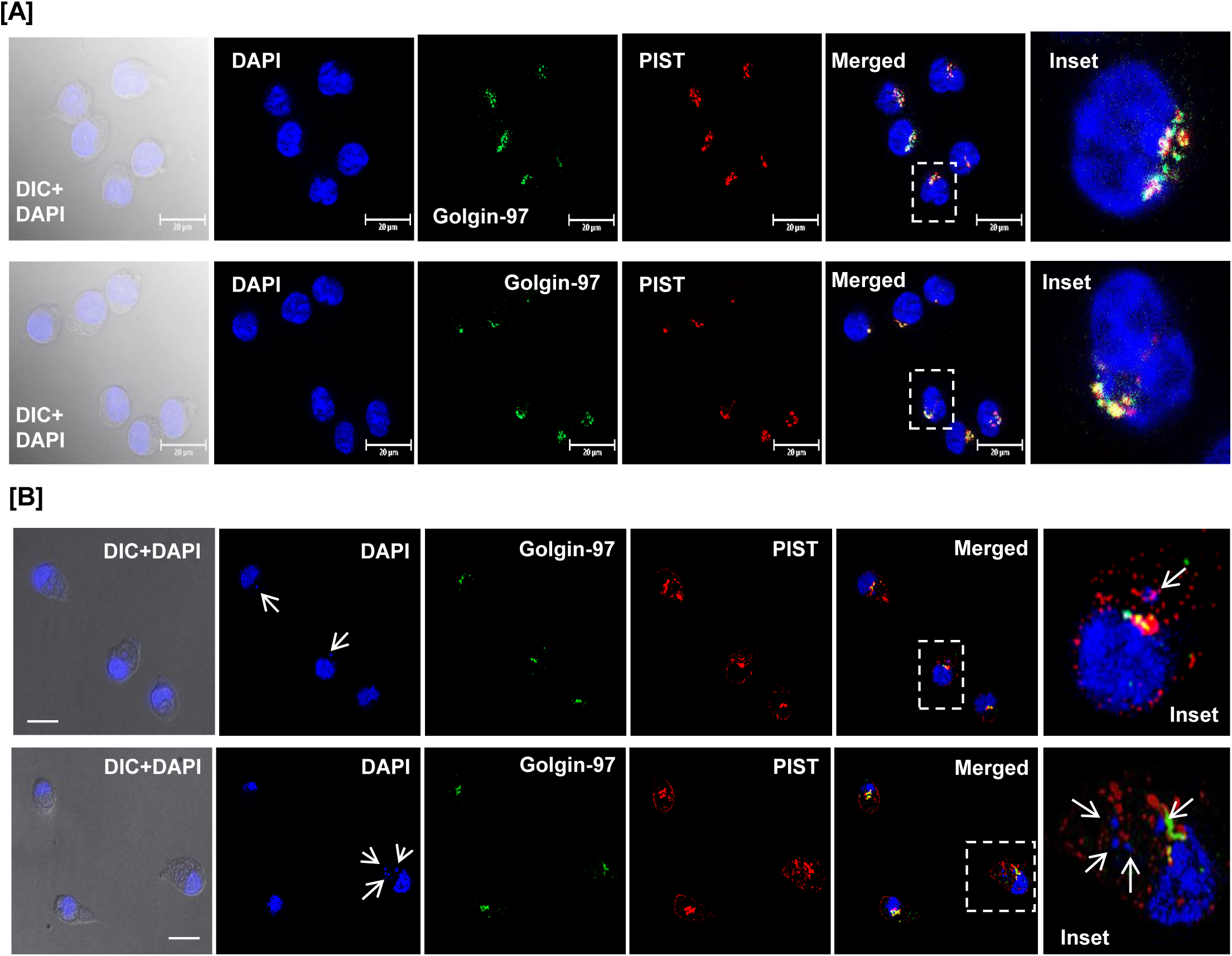
*L. major* infection alters macrophage PIST localization from Golgi compartment. (A) J774A.1 macrophage cells were immunostained with anti-Golgin-97 (Golgi marker) (green) and anti-PIST (red) and further the colocalization status between these two proteins was monitored. Nuclei was stained with DAPI. Under resting condition within J774A.1 macrophage cells PIST completely colocalized with Golgin-97. (B) Balb/c isolated peritoneal macrophage cells were first infected with *L. major* promastigotes for 6hrs. Cells were then fixed and immunostained similarly as mentioned above. Arrows in DAPI stained panel indicative of parasite nuclei. Inset panel shows the altered distribution of PIST within Lm infected peritoneal macrophages where the protein is present within parasite containing compartment. Both (A) and (B) shows two different representative fields of the cells. Cells were visualised with Zeiss Apotome microscope using ×100 oil immersion objective. Scale bar: 20μm.

**Figure S3.**
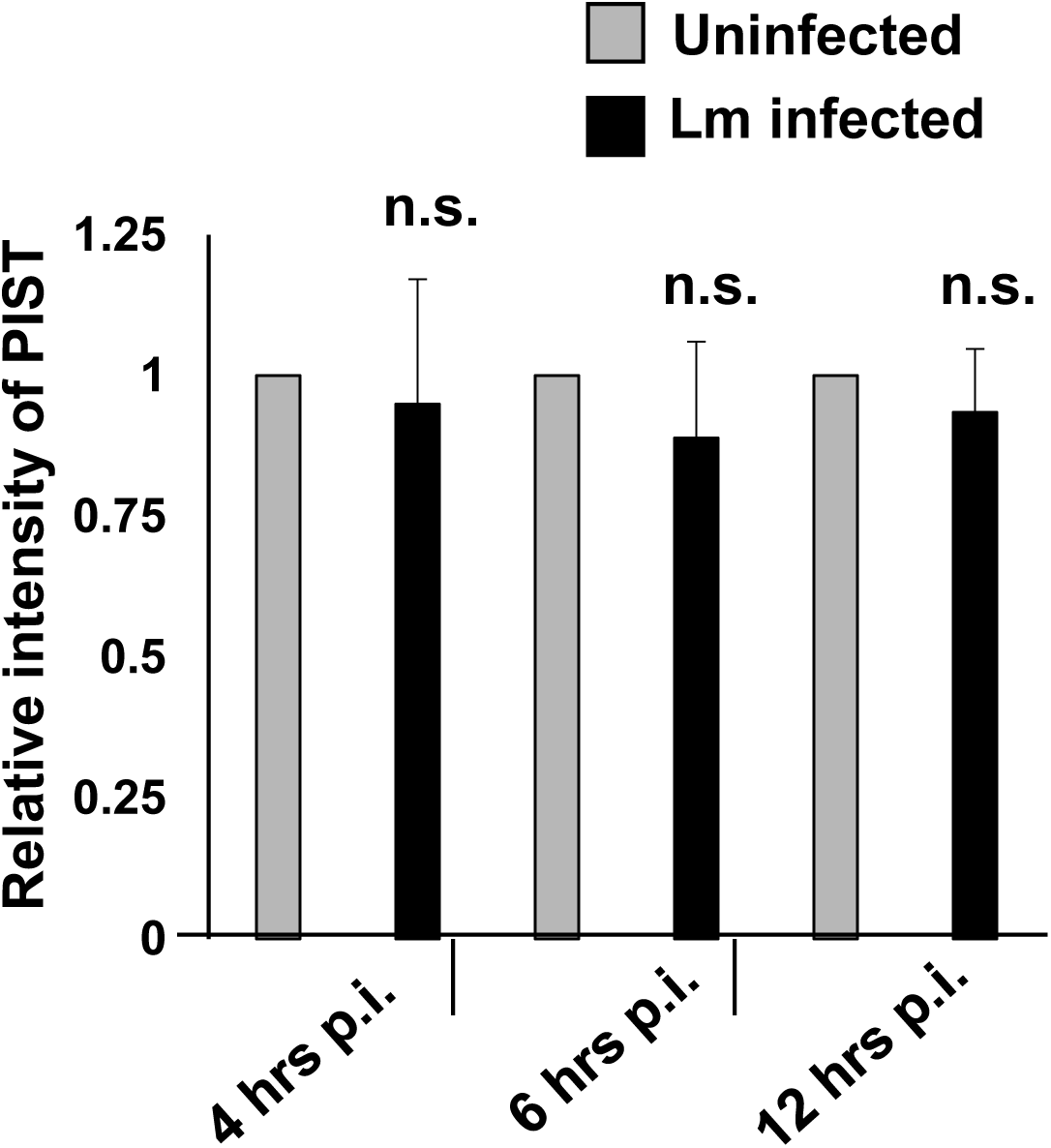
Quantification of macrophage PIST protein level during *L. major* infection by western blot analysis. J774A.1 macrophage cells were either uninfected or infected with *L. major* promastigotes for 4- 12 hrs. After 4, 6 and 12 hrs of infection cell lysates were prepared from both uninfected and Lm infected macrophage cells and these were subjected to western blot assay using anti-PIST. GAPDH was used as the loading control. The relative intensity of PIST was measured using ImageJ software against GAPDH from three different experimental sets and represented here as the bar diagram where grey bar represents uninfected cells and black bar represents Lm infected macrophage cells. Error bars represent Mean± SD, values calculated from at least three independent experiments. n.s. non-significant.

**Figure S4.**
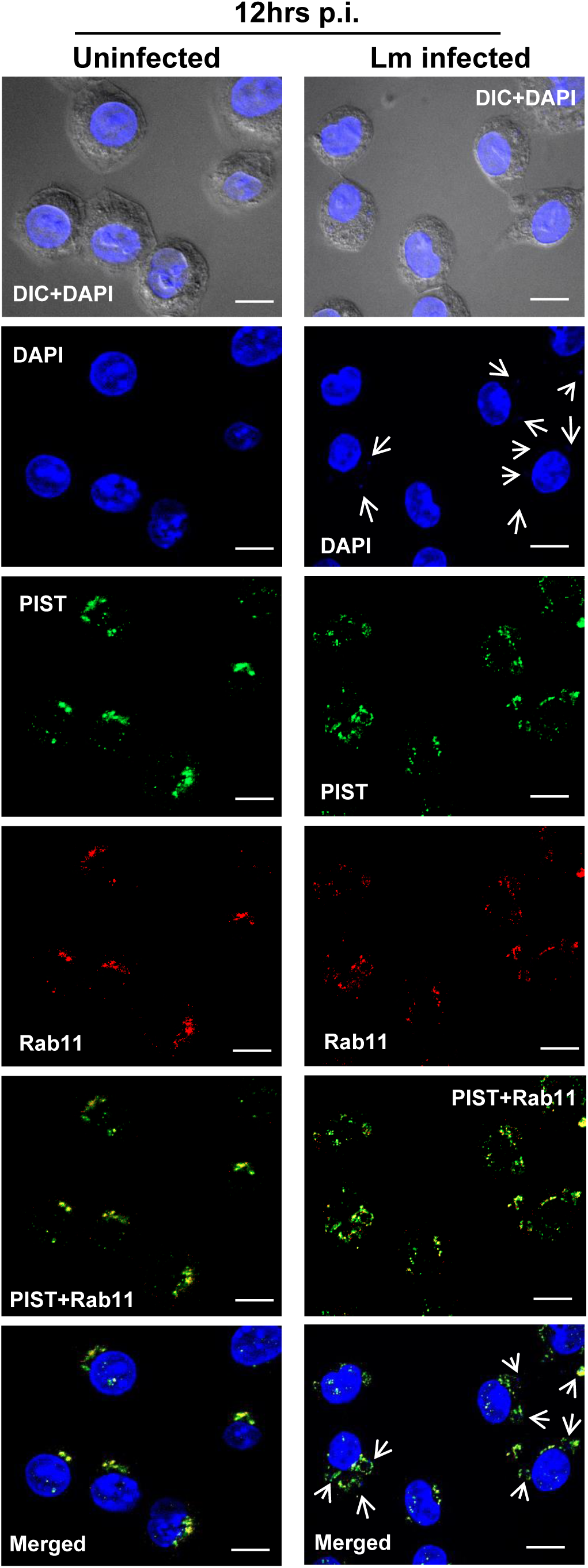
PIST colocalizes with Rab11 in *L. major* infected macrophages. J774A.1 macrophage cells either uninfected or infected with *L. major* promastigotes for 12hrs and further the cells were fixed and immunostained with anti-PIST (green) and anti-Rab11 (red). The colocalization status between these two proteins was monitored. Nuclei was stained with DAPI. Arrows in DAPI stained panel indicative of parasite nuclei. Within Lm infected macrophages PIST is highly colocalized with Rab11. Both uninfected and Lm infected panels are showing representative fields of the cells. Cells were visualised with Zeiss Apotome microscope using ×63 oil immersion objective. Scale bar: 20μm.

**Figure S5.**
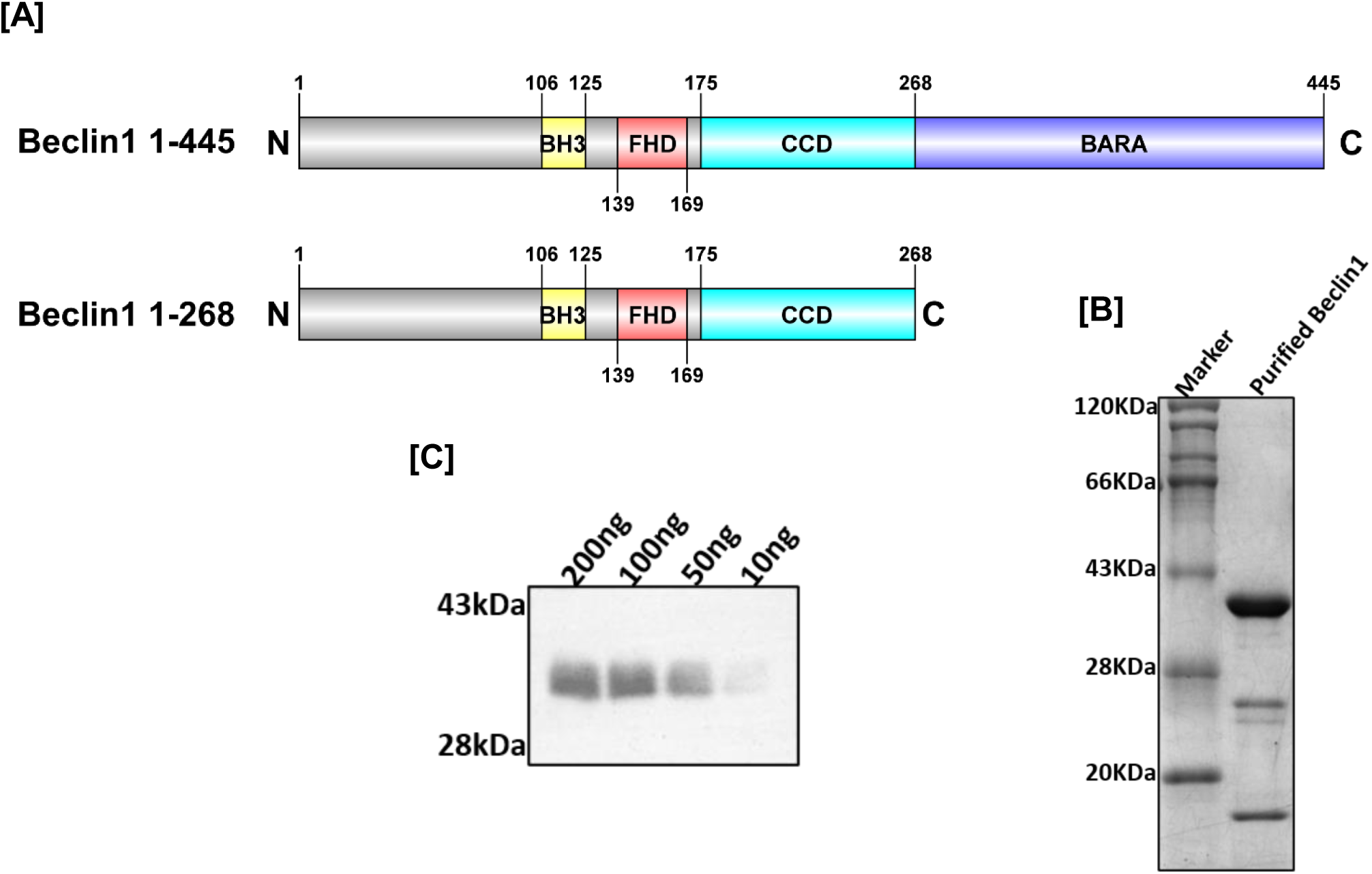
Generation of anti- Beclin 1 polyclonal antiserum. (A) Schematic diagram of mouse Beclin 1 (1-445) and mouse Beclin 1 fragment 1-268 construct against which the antiserum was generated. Beclin 1 contains a BCl-2 homology domain 3 (BH3 motif; 106-125), a flexible helical domain (FHD; 139-169), a coiled-coil domain (CCD; 175-268) and a β-α repeated autophagy domain (BARAD; 268-445), Beclin 1 1-268 fragment construct lacking this BARA domain, the structure of Beclin 1 and its domains were created using software DOG2.0. (Ren et al., 2009). (B) Purified C-terminal Histidine tagged recombinant Beclin 1 expressed into *E. coli* BL-21 DE3 strain used to raise rabbit anti-beclin1 polyclonal antibody. (C) Different concentration of recombinant Beclin 1 1-268 fragment (10ng, 50ng, 100ng, and 200ng) were analysed by western blot with immunized rabbit serum at dilutions of 1:1000.

**Figure S6.**
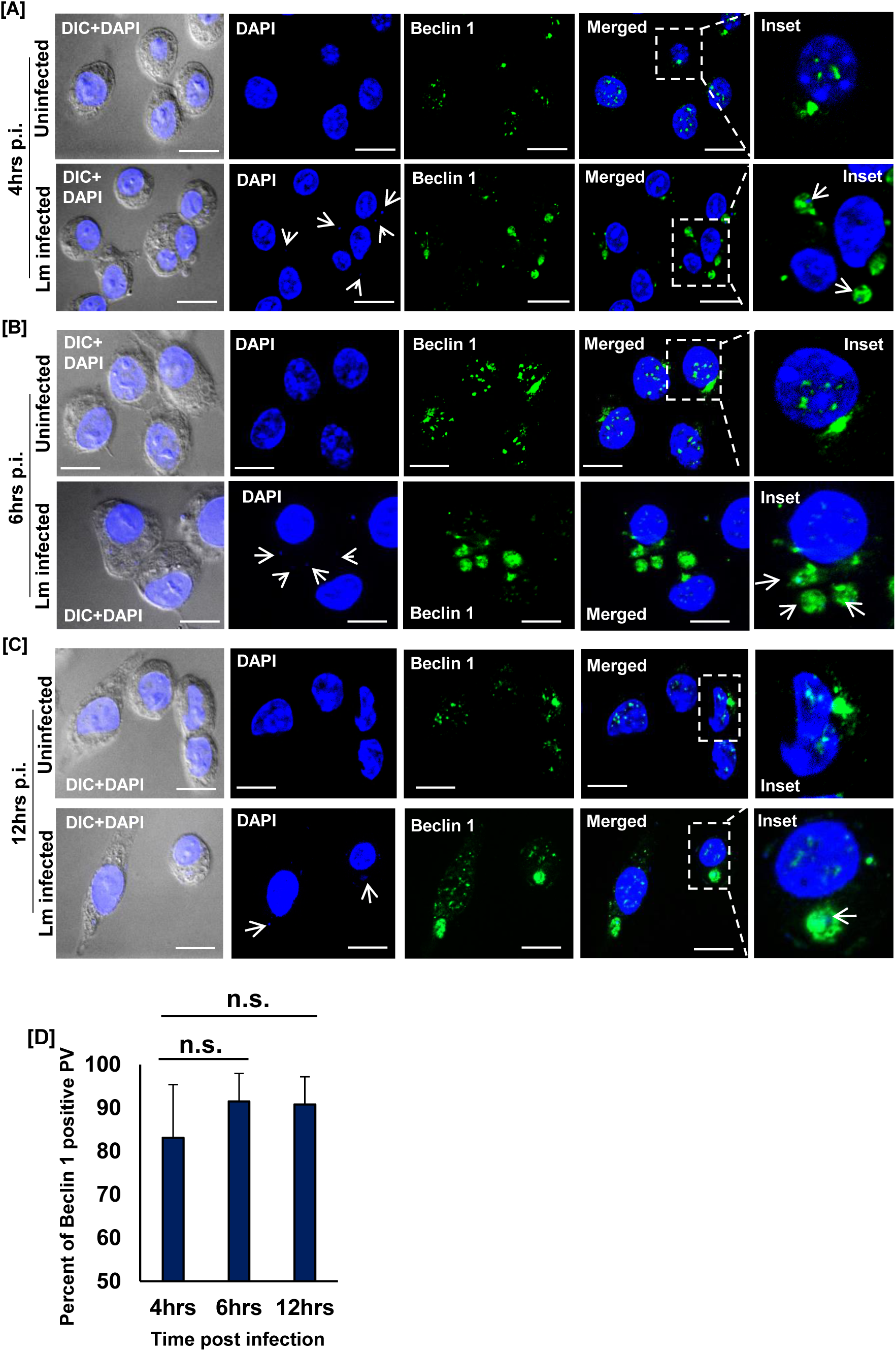
Beclin 1 is recruited to the parasite containing compartment in infected macrophages. J774A.1 macrophages were infected with *L. major* (Lm) promastigotes and immunostained with anti-Beclin 1 (green) at (A) 4 hrs, (B) 6 hrs and (C) 12 hrs post infection. Nuclei were stained with DAPI (blue). (DIC+DAPI) panel in each time points shows the uninfected and Lm infected macrophage cells where in the DAPI panel the presence of intracellular parasites (smaller nuclei) as indicated by white arrows within infected cells. Insets shows magnified image of a representative cell from both uninfected and Lm infected panels with distribution of Beclin 1 protein and arrows indicating parasite nuclei. Cells were visualised with Zeiss Apotome microscope using ×63 oil immersion objective. Scale bar: 20μm. (D) Bar diagram represents percent of Beclin 1 positive parasitophorous vacuole (PV) at 4- 12 hrs p.i. At least 20 cells from three independent experiments were scored in each of the cases. Error bars represent mean± standard deviation calculated from at least three independent experiments. n.s. non-significant.

**Figure S7.**
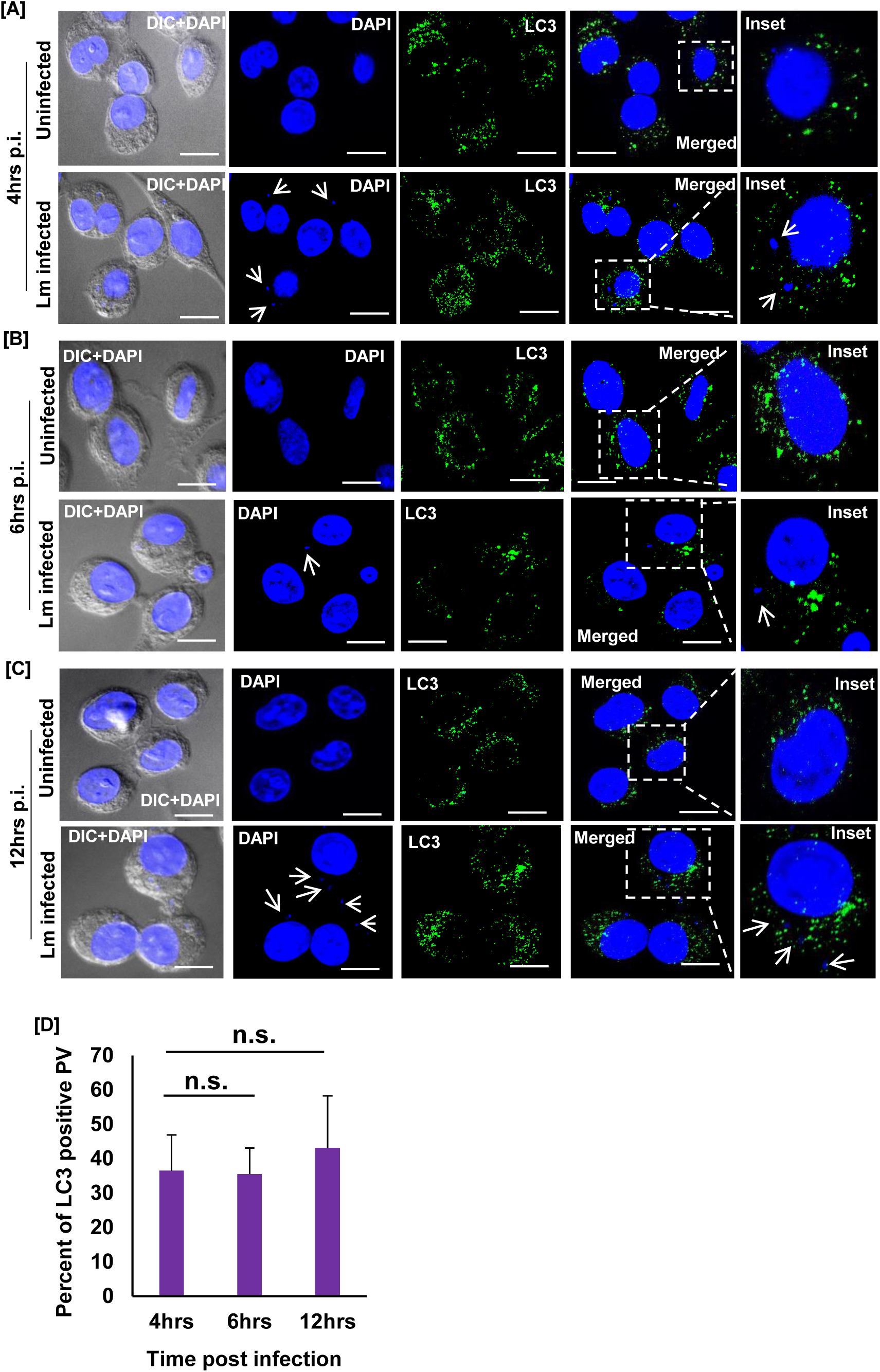
Parasite containing compartment is devoid of LC3 in *L. major* infected macrophages. J774A.1 macrophages were infected with *L. major* (Lm) promastigotes and immunostained with anti-LC3 (green) at (A) 4 hrs, (B) 6 hrs and (C) 12 hrs post infection. Nuclei were stained with DAPI (blue). (DIC+DAPI) panel in each time points shows the uninfected and Lm infected macrophage cells where in the DAPI panel the presence of intracellular parasites (smaller nuclei) as indicated by white arrows within infected cells. Insets shows magnified image of a representative cell from both uninfected and Lm infected panels with distribution of LC3 protein and arrows indicating parasite nuclei. Cells were visualised with Zeiss Apotome microscope using ×63 oil immersion objective. Scale bar: 20μm. (D) Bar diagram represents percent of LC3 positive parasitophorous vacuole (PV) at 4- 12 hrs p.i. At least 20 cells from three independent experiments were scored in each of the cases. Error bars represent mean± standard deviation calculated from at least three independent experiments. n.s. non-significant.

